# While shoot herbivory mitigates, root herbivory exacerbates eutrophication’s impact on diversity in a grassland model

**DOI:** 10.1101/799528

**Authors:** Michael Crawford, Ulrike E. Schlägel, Felix May, Susanne Wurst, Volker Grimm, Florian Jeltsch

## Abstract

Eutrophication is widespread throughout grassland systems and expected to increase during the Anthropocene. Trophic interactions, like aboveground herbivory, have been shown to mitigate its effect on plant diversity. Belowground herbivory may also impact these habitats’ response to eutrophication, but the direction of its influence is much less understood, and likely to depend on factors such as the herbivores’ preference for dominant species and the symmetry of belowground competition. If preferential towards the dominant, fastest growing species, root herbivores may reduce these species’ relative fitness and support diversity during eutrophication. However, as plant competition belowground is commonly considered to be symmetric, root herbivores may be less impactful than shoot herbivores because they do not reduce any competitive asymmetry between the dominant and subordinate plants.

To better understand this system, we used an established, two-layer, grassland community model to run a full-factorially designed simulation experiment, crossing the complete removal of aboveground herbivores and belowground herbivores with eutrophication. After 100 years of simulation, we analyzed communities’ diversity, competition on the individual-level, as well as their resistance and recovery. The model reproduced both observed general effects of eutrophication in grasslands and the short-term trends of specific experiments. We found that belowground herbivores exacerbate the negative influence of eutrophication on Shannon diversity within our model grasslands, while aboveground herbivores mitigate its effect. Indeed, data on individuals’ above- and belowground resource uptake reveals that root herbivory reduces resource limitation belowground. As with eutrophication, this shifts competition aboveground. Since shoot competition is asymmetric—with larger, taller individuals gathering disproportionate resources compared to their smaller, shorter counterparts—this shift promotes the exclusion of the smallest species. While increasing the root herbivores’ preferences towards dominant species lessens their negative impact, at best they are only mildly advantageous, and they do very little reduce the negative consequences of eutrophication. Because our model’s belowground competition is symmetric, we hypothesize that root herbivores may be beneficial when root competition is asymmetric. Future research into belowground herbivory should account for the nature of competition belowground to better understand the herbivores’ true influence.

## 2 Introduction

Eutrophication, i.e. excessive nutrient (e.g. N and P) deposition into ecosystems, reduces diversity worldwide and may likely worsen as globalization drives land-use intensification (Vitousek et al. 1997; Lambin et al. 2001; Steffen et al. 2015). In terrestrial ecosystems, grasslands are heavily impacted by eutrophication (Stevens et al. 2004; Dupré et al. 2009; Bobbink et al. 2010; Hautier et al. 2014). Those in central Europe, for example, have seen reductions in plant diversity of 50% over the last 50 years, mostly attributable to local nutrient input and land use intensification (Wesche et al 2012).

These negative impacts emerge as competitive plant functional types (PFTs) begin to dominate the community (Harpole and Tilman 2007; Hautier et al. 2009). No longer resource-limited, these species—well adapted to quickly converting nutrients into biomass—rapidly grow to overshadow their smaller neighbors. Their initial superiority is further compounded by the asymmetry inherent in aboveground competition, with taller plants obtaining disproportionately more light than their shorter counterparts (Weiner et al. 1986; DeMalach et al. 2016, 2017; Hautier et al. 2018). This asymmetry is the primary reason trophic interactions are recognized as an important mechanism through which grassland diversity can resist and potentially recover from eutrophication.

Aboveground herbivory tends to inhibit competitive exclusion by reducing the competitive asymmetry between the largest and smallest plants. By disproportionately affecting the largest, fastest growing functional types, aboveground herbivory increases the light available to smaller individuals (Borer et al. 2014). Thus, aboveground herbivory presents a countervailing force that may constrain the species loss due to eutrophication, safeguarding diversity by decreasing the performance of the strongest species (Olff and Ritchie 1998; Anderson et al. 2018; Mortensen et al. 2018; but see Borgström et al. 2016). Further, several recent studies have also indicated that trophic interactions such as herbivory can increase the resilience of grasslands to stress through trophic compensation (Thébault and Fontaine 2010; Eisenhauer et al. 2011; Giulia et al. 2015), though not eutrophication *per se*.

While the role of aboveground herbivory in mitigating the impact of eutrophication on diversity has been established, the impact of its counterpart belowground is poorly understood (Blossey and Hunt-Joshi 2003). Despite 40–70% of annual net primary production being belowground (Vogt et al. 1995) and root herbivores likely removing as much, or more, biomass than their foliar cousins (Zvereva and Kozlov 2012; Kozlov and Zvereva 2017), practical obstacles in its research have left belowground herbivory historically “out of sight, out of mind,” (Hunter 2001). The two studies that have investigated root herbivory’s role in the response of grassland systems to eutrophication have shown that herbivores compound its effect, further decreasing biodiversity (La Pierre et al. 2014; Borgström et al. 2017).

Most recently, Borgström et al. (2017) designed a factorial experiment, crossing the presence of aboveground herbivores, belowground herbivores, and eutrophication. They found that while aboveground herbivory decreased the relative biomass of grasses and thus counteracted the impact of eutrophication, root herbivores consistently decreased diversity, regardless of the other treatments. These results are striking: Above- and belowground herbivory are not equivalent. This highlights the need to investigate the underlying mechanisms behind this difference more deeply. By further developing this understanding, we will improve our ability to predict biodiversity in grassland ecosystems, as well as their response to eutrophication.

Some evidence shows belowground herbivores are preferential towards larger root systems, rather than generalist, and that they may reduce these plants’ dominance and support diversity (Sonnemann et al. 2012, 2015). Indeed, several studies have shown that belowground herbivores increase diversity, albeit in non-eutrophic systems (De Deyn et al. 2001; Stein et al. 2010). However, the mechanisms underlying above- and belowground herbivory may be more complicated than comparing their feeding preferences alone. This is because contrary to plant competition aboveground, belowground resources are most often symmetrically allocated based on plant size (Schwinning and Weiner 1998). Without a competitive asymmetry to equalize, root herbivores are unlikely to foster diversity maintenance (Chesson 2000). By way of analogy, if aboveground competition were not size-asymmetric, size asymmetric herbivory would be less effective in increasing the community’s diversity.

To summarize, the literature suggests that belowground herbivores do not necessarily mitigate eutrophication, but their effects may hinge on their preference towards competitive species as well as the symmetry of belowground competition. To break down how root herbivory influences the diversity and resilience of grassland systems—and their reaction to eutrophication—it would be helpful to test in isolation not just the effect of herbivory (above- and belowground) and eutrophication, but also of the belowground herbivores’ preferences towards dominant PFTs. These nuances are ripe to be investigated with ecological modeling, which enables researchers to continuously monitor high-resolution variables describing not only the plants’ state but also their above- and belowground competitive environment. Further, given that the full extent of eutrophication’s influence may only emerge over the long-term (Kidd et al. 2017), modeling may provide additional useful insights.

In this paper, we extend on the empirical short-term results of Borgström et al. (2017) by implementing their factorial design inside of an established grassland community model. We then parameterize the feeding preferences of the belowground herbivores to reflect a gradient from generalist to preferential. While “generalists” will consume all species equally, proportional to their biomass, “preferential” herbivores will disproportionately focus on the dominant species within the grassland, asymmetrically consuming those species that have larger root systems.

We also expand on their work by examining herbivores’ impact on the resilience of grasslands to eutrophication. Despite some studies predicting that trophic interactions will increase the stability of ecological systems (Thébault and Fontaine 2010; Eisenhauer et al. 2011; Giulia et al. 2015), no studies have examined how belowground herbivores mediate the response of grasslands to stresses like eutrophication. Therefore, in addition to examining the impact of herbivores on grasslands diversity *per se*, we also investigate how the removal of herbivores (above- and belowground) impact the resistance and recovery (sensu Hodgson et al. 2015) of these grasslands to eutrophication.

## 3 Methods

We used the individual-based and trait-based, spatially-explicit grassland assembly model, IBC-grass (May et al. 2009), that incorporates above- and belowground herbivory. Since its introduction (May et al. 2009), IBC-grass has been used to investigate numerous aspects of grassland dynamics, from resilience (Weiss and Jeltsch 2015; Radchuck et al. 2019), to species coexistence (Pfestorf et al. 2016; Crawford et al. 2018), and ecotoxicology (Reeg et al. 2017; Reeg et al. 2018). Importantly, Weiss et al. (2014) parameterized the model with data from trait databases and a survey of plant functional types from German grasslands collected through the Biodiversity Exploratories (Fischer et al. 2010; Pfestorf et al. 2013). With this empirical parameterization the model successfully reproduces, without calibration at the community level, empirically-observed grassland biodiversity patterns (Weiss et al. 2014).

A full description of the model can be found in the ODD (Overview, Design concepts, Details) protocol (Grimm et al. 2010, 2006; Appendix S1). The following is a summary of the model as well as an explanation of our modifications to it.

### 3.1 Overview of the IBC-grass model

IBC-grass simulates local community dynamics on a 141 x 141 cell grid, where each cell corresponds to 1 cm^2^ (resulting in a roughly 2 m^2^ grid space) and can hold one plant’s stem. Its time step corresponds to one week, and there are 30 weeks per year representing the vegetation period. A plant is characterized by its functional traits and the biomass of its three distinct compartments: aboveground mass, belowground mass, and reproductive mass.

A plant’s competitive area is defined by an aboveground and a belowground “zone of influence” (ZOI). The two ZOIs are both circular areas around the plant’s stem, from which it acquires either above-or belowground resources (Schwinning and Weiner 1998; Weiner et al. 2001). While the plant’s stem is contained within one grid cell, its ZOI can cover many. Belowground, the area of a plant’s ZOI is related directly to its root biomass (Appendix S1: Eq. A3a). Aboveground, the area is the product of its aboveground biomass as well as two functional traits, leaf-mass ratio (LMR) and specific leaf area (SLA) (Appendix S1: Eq. A1). The plant’s LMR describes its proportion of photosynthetically active (leaf) tissue to the total shoot tissue and its SLA is a constant ratio between leaf mass and ZOI area. In IBC-grass, a plant with a low LMR will generally have a small shoot area, but overshadow its shorter neighbors; a high SLA corresponds to a leaf that is larger and therefore able to gather more aboveground resources, but also less well defended from aboveground herbivory than its lower-SLA counterparts.

When two plants’ ZOIs overlap they compete for resources. A cell’s aboveground resources correspond to light while its belowground resources—likewise unidimensional—therefore correspond to water and nutrients. The proportion of a cell’s resources a plant obtains during competition depends on its competitive abilities and how many other ZOIs overlap the cell. Aboveground competition is size-asymmetric, i.e. the larger plant takes up resources from each contested cell not only in proportion to its competitive ability (measured as the maximum units it can acquire per week, termed its g_max_), but also in proportion to its aboveground mass and LMR^-1^, reflecting its height advantage over the smaller plants (Appendix S1: Eq. A3b). In other words, aboveground competition disproportionately favors the larger, taller competitor. Belowground competition, however, is size-symmetric, i.e. only their competitive abilities (their g_max_) are considered (Appendix S1: Eq. A3a).

Intraspecific competition is also included in the form of negative-density dependent competition, reflecting species-specific predators or pathogens (May et al. 2009). This density-dependent competition was modeled by decreasing the resource uptake of an individual in proportion to the square-root of the number of conspecifics in its neighborhood ZOI (Appendix S1: Eq. A3c).

All grid cells’ total resources are kept constant through space and time; only a plant’s biotic neighborhood influences the amount of resources available to it at any given time step. When a plant is unable to gather enough resources, it changes its resource allocation between above- and belowground parts, depending on which compartment is more limited (i.e. shoot or root). If resource uptake in either of the two compartments is below a certain threshold, the plant is considered stressed. Each consecutive week a plant is stressed increases its chance of mortality linearly, in addition to a background, stochastic, annual mortality of 20%.

The plants are characterized by four sets, or syndromes, of functional traits, a subset of those proposed as the “common core list of plant traits” by Weiher et al. (1999). The first set of traits defines the plant’s maximum size (m_max_), which positively correlates with seed size and negatively correlates with dispersal distance (Eriksson and Jakobsson 1998; Jongejans and Schippers 1999). The second defines the plant’s growth form, or leaf to mass ratio (LMR), which describes the plant as either a rosette, erect, or intermediate growth form type. The third set defines its competitive ability, or maximum resource utilization per time step (g_max_), and negatively correlates with its stress tolerance (Grime 2001). The fourth trait set describes the plant’s grazing response, positively correlating its palatability with its specific leaf area (SLA) (Westoby et al. 2002).

### 3.2 Aboveground herbivory

Aboveground grazing events, modelling the herbivory of large mammals, reflect the partial removal of a plant’s aboveground biomass. The frequency of grazing is specified by a constant weekly probability (p_graz_) of a grazing event. The grazers tend to act selectively towards certain traits, with a preference for larger, taller individuals exhibiting high SLAs, which corresponds to relatively palatable leaves (Díaz et al. 2001).

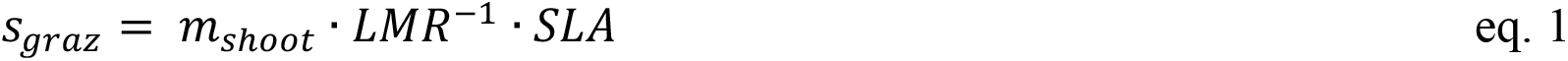

The probability for a given individual to be grazed within one week is derived as its grazing susceptibility (s_graz_) in proportion to the current maximum individual susceptibility of all the plants (in other words, the susceptibility of the most-susceptible plant). All plants are checked to be grazed in a random order. If a plant is grazed, 50% of its shoot mass and all of its reproductive mass are removed. The random choice of plants is repeated without replacement until 50% of the total aboveground biomass on the grid has been removed or the residual biomass is reduced to less than (15 g/m², Schwinning and Parsons 1999) what is considered ungrazable. After a pass through the entire plant community, if either of these two end conditions are unmet the process is repeated. This allows a plant individual to be grazed never or several times during one week with a grazing event.

In this study, we use a grazing probability of 0.2 per time step, consistent with previous studies using this submodel.

### 3.3 Belowground herbivory

Belowground herbivory was implemented such that each time step some percentage of the extant biomass is removed from each of the plants, with a gradient of preference in root size ranging from generalist to preferential (i.e. disproportionally eating larger root systems). This herbivory algorithm is intended to reflect the influence of belowground, invertebrate herbivores, such as those belonging to the genus *Agriotes*, one of the most abundant root herbivores in Europe. As this genus generally tends to eat plants with high biomass and growth rates (Sonnemann et al. 2012, 2015), we refrain from explicitly modelling the plants’ roots palatability.

The feeding need at week 𝑡, 𝑛*_t_*, is calculated as a defined percent (feeding rate, 𝑓) of that week’s expected root mass, which is estimated by averaging each previous week’s total realized root biomass 𝑅*_i_* for the previous 𝑤 weeks,

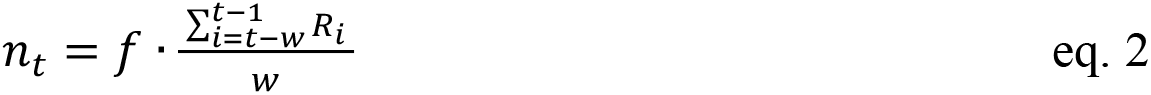

For this analysis, the feeding rate (𝑓) is 0.1 per week, potentially lower than the typical belowground herbivory pressure (Zvereva and Kozlov 2012), but equal to the aboveground herbivory pressure commonly used in IBC-grass. The number of weeks used to estimate the expected root mass, 𝑤, is 10. Both parameters are held constant in the following analysis.

The biomass to be removed from each individual’s root mass (𝑔*_i,t_*) is calculated each week as:

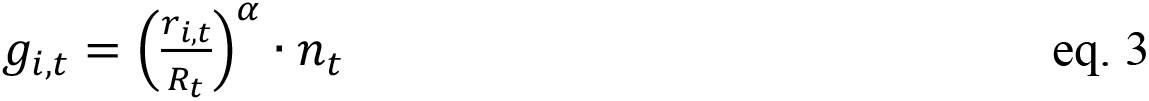

where 𝑟*_i,t_* is the expected root mass of individual 𝑖 in week 𝑡 and 𝑅*_t_* is the week’s realized total root mass, which may differ from the expected root mass (Appendix S1: Fig. A3). 𝑅*_t_* differs from 𝑅*_i_* in eq. 2 in that 𝑅*_t_* refers to the total realized root biomass on week 𝑖 (and ranging backwards by 𝑤 weeks), whereas 𝑅*_t_* refers to the total realized root biomass on the current week. The parameter 𝛼 represents the generality of the herbivory; set at 𝛼 = 1, 𝑔*_i,t_* will equal the plant’s root mass (𝑟*_i,t_*) in proportion to the total root mass (𝑅*_t_*) at time 𝑡. Above 1, 𝛼 will increase the preference of the herbivores to disproportionally prefer large root systems (Sonnemann et al. 2013). This parameter is varied from 1 (generalist) to 2 (extremely preferential). If the biomass to be removed from a plant is larger than its total root mass (which may occur, based on the distribution of plant biomasses and 𝛼), the plant is killed and the overshoot biomass remains in the feeding need (𝑛*_t_*), to be removed from other plants.

### 3.4 Eutrophication

Eutrophication was simulated as an increase in belowground resources (BRes) from the baseline resource rate. Therefore, in IBC-grass a eutrophication intensity of 10 would translate to an increase in belowground resources of 10 BRes for the duration of the experiment. Immediately after the experimental period, the belowground resources return to their pre-eutrophication level for 100 simulation years, for the analysis of the plots’ recovery. Although abiotic modifications to natural communities (e.g. nutrient, herbicide, or pesticide addition) will degrade more slowly than is modelled in our present work, we argue that as a first approximation, this simplifying assumption will demonstrate—in principle—how the biotic community will respond to the cessation of these human-caused disturbances. For this analysis we increased the amount of belowground resources by 50% over their baseline levels (Weiss et al. 2014), increasing from 60 to 90 BRes during the experimental period.

### 3.5 Simulation design and experiments

We implement a full-factorial design mirroring Borgström et al. (2017) inside of IBC-grass. After a burn-in period of 100 simulation-years wherein the communities settle into quasi-equilibrium, they are experimentally manipulated through the application of aboveground herbivore exclusion, belowground herbivore exclusion, and eutrophication, fully crossed. Before the experimental treatments begin, all simulations have a moderately-low level of baseline belowground resources (60 resource units) and both above- and belowground herbivory. The belowground herbivory is parameterized along a gradient of preferentiality, such that each community has one of five levels from 1 (generalist) to 2 (very preferential, see *Methods: Belowground Herbivory*). During the experimental period, aboveground and belowground herbivore exclusion is modelled as the complete elimination of these two submodels. This period lasts for 100 simulation years, long enough for all communities to reach a quasi-equilibrium once again. After the experimental window ends, the presence of above- and belowground herbivores, as well as the level of belowground resources returns to their pre-experiment values and the simulation is left to run for another 100 years, so that the communities’ recovery can likewise be examined. Each parameterization is replicated 50 times.

To understand how these three factors (above- and belowground herbivory, and eutrophication) influence the diversity of our model grasslands, we examine the simulated communities’ Shannon diversity—which combines the effects of richness and evenness—shortly after the experiment begins and immediately before it concludes. We then investigate the corresponding shifts in the individuals’ resource uptake levels (a proxy for competition). To understand the grasslands’ resilience dynamics, we also inspect two key resilience metrics (*sensu* Hodgson et al. 2015), resistance and recovery. Resistance is the magnitude of change some metric (e.g. diversity) undergoes directly after a disturbance. In our case, we were exploring the effects of eutrophication on diversity with and without above- and belowground herbivory. Experimentally removing the herbivores was thus a diagnostic—or proximate disturbance— meant to reveal the stabilizing effects of herbivory, while the ultimate disturbance of interest was eutrophication. Recovery is the amount of time needed for a system to reach its original state (measured as time to recovery, TTR). Here we explored recovery, with and without eutrophication, after herbivory was reinstated. Recovery necessitates some external seed input, so for the duration of the recovery period we add a modicum of seeds—*seeds* 𝑚^-2^ *PFT*^-2^ *year*^-1^ (Weiss et al. 2014, Reeg et al. 2018)—to reintroduce any species extirpated during the experimental phase. A community is considered to have recovered once its diversity returns to within two standard deviations of the control communities’—no herbivory removal or eutrophication—diversity.

## 4 Results

After 100 years, both eutrophication and the presence of generalist belowground herbivory decrease Shannon diversity, while aboveground herbivory increases it (Fig. 1). A linear model predicting Shannon diversity through the three-way interaction of these variables (Table 1) revealed significant interactions between eutrophication and both above- and belowground herbivory. While aboveground herbivory can partly mitigate the negative impacts of eutrophication on diversity, belowground herbivory exacerbates it. Interestingly, while there was no interaction between above- and belowground herbivory at ambient levels of belowground resources, a significant three-way interaction with eutrophication indicates that—in grasslands with eutrophication—combined above- and belowground herbivory result in a significantly lower Shannon diversity than if there was no interaction present (i.e. their effects were additive). This interaction was larger than the negative effect of belowground alone.

**Figure 1.**
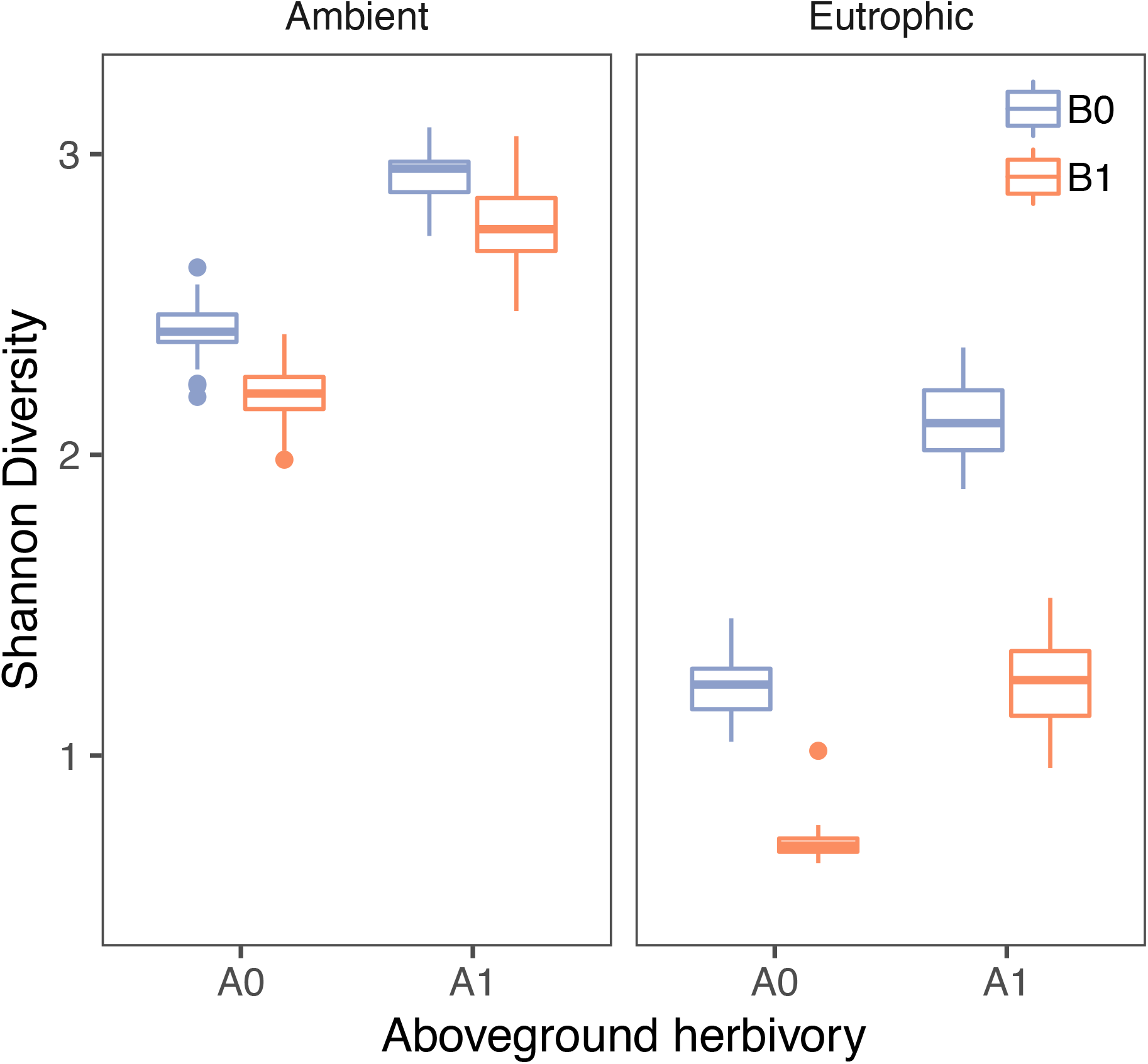
Shannon diversity after 100 years of experimental treatment. Control treatment retains above- and belowground herbivory with ambient resource levels. A0 – Aboveground herbivory removed, A1 – Aboveground herbivory present; B0 – Belowground herbivory removed, B1 – Belowground herbivory present. While aboveground herbivory increases diversity, belowground herbivory has a negative effect on diversity, exacerbating the negative effect of eutrophication.

**Table 1:** Impact of above- and belowground herbivory on the diversity response of simulated grassland plots to eutrophication.

| Variable | Estimate | Std. Error | t-value | p-value |
| --- | --- | --- | --- | --- |
| Intercept | 2.413 | 0.015 | 164.669 | < **0.001** |
| Eutrophication | -1.177 | 0.021 | -56.783 | < **0.001** |
| A1 | 0.522 | 0.021 | 25.188 | < **0.001** |
| B1 | -0.212 | 0.021 | -10.240 | < **0.001** |
| Eutrophication:A1 | 0.357 | 0.029 | 12.197 | < **0.001** |
| Eutrophication:B1 | -0.319 | 0.029 | -10.879 | < **0.001** |
| A1:B1 | 0.024 | 0.029 | 0.831 | 0.407 |
| Eutrophication:A1:B1 | -0.36 | 0.041 | -8.894 | < **0.001** |
| Adjusted R-squared: 0.981; F-statistic: 2976 on 7 and 392 DF |  |  |  |  |

Comparing our results to Borgström et al. (2017), the patterns of Pielou’s evenness (E) shortly after the experimental period begins (3 years, mirroring Borgström et al. 2017) are broadly concordant (Appendix S2: Fig. 1).

### 4.1 Variation with root herbivore preference

We next shift our attention from generalist belowground herbivores to those with increasing preference towards dominant species, by testing a gradient from generalist herbivores (𝛼 = 1) to those extremely preferential towards large root systems (𝛼 = 2). We found that increasing the herbivore’s preference towards dominant plants increased diversity relative to purely generalist herbivores, but that this effect was insufficient to mitigate the negative effects of eutrophication (Fig. 2). Although in grasslands with ambient resources, very preferential herbivores had no impact on diversity (especially compared to generalist herbivores, which significantly reduced it), eutrophication reduced diversity by threefold the positive impact of preferential herbivores, overshadowing any positive influence of preferential herbivory. Therefore, preferential herbivores, at most, neutrally impact diversity.

**Figure 2.**
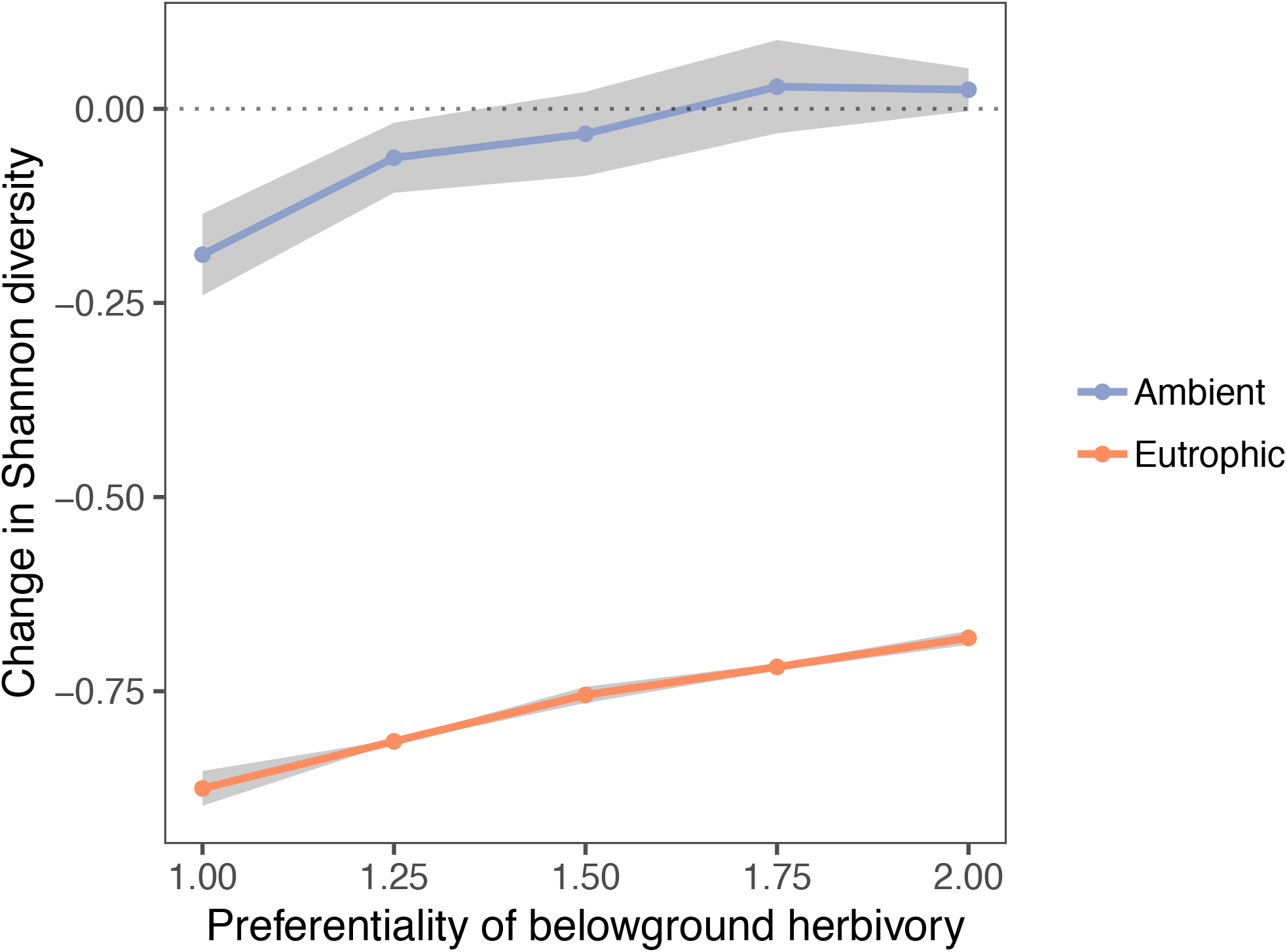
Change in Shannon diversity (relative to the diversity with belowground herbivores excluded) with increasing degree of belowground herbivore preference towards the largest plant functional types. Aboveground herbivory is held constant and present. Ribbons indicate one standard deviation around the mean. Preferentiality of root herbivores only slightly lessens the negative effect of root herbivory on diversity. However, this is not sufficient to mitigate the negative influence of eutrophication.

To understand how preferential herbivory differentially impacted species fitness more deeply, we isolated the dominant PFT from each community with generalist herbivores, defining it to be the PFT with the largest summed root biomass. We then plotted its total root biomass under the gradient of herbivore selectivity, and found that only under ambient belowground resource levels do higher levels of selectivity decrease the total root biomass of the most dominant PFTs in the community (Appendix S2: Fig. 2). Under eutrophication, there is no interaction between that PFT’s root biomass and the herbivores’ preference for larger root systems.

### 4.2 Competition on the individuals’ level

We also measured the shoot- and root- resource uptake per individual after the experimental period. This measure, the ratio of resource uptake to biomass above- or belowground, is a proxy for competition in each compartment; a high ratio reflects less competition, because the plant is receiving more resources per unit biomass. Lower values, therefore, reflect higher competition wherein few resources are available for the individual to take up. The average ratio between an individual’s aboveground resources uptake to it shoot biomass (𝐴𝑅𝑒𝑠 ∶ 𝑚*_shoot_*) increased in the presence of aboveground herbivores (Fig. 3A). This reflects a decrease in aboveground competition, as the removal of aboveground biomass reduces the number of plants competing for each cells’ resources. The introduction of belowground herbivory mitigated this impact, in effect reducing aboveground herbivory’s ability to decrease aboveground competition. Eutrophication was even more disruptive to the efficacy of aboveground herbivory, drastically reducing the aboveground resource uptake ratio. This suggests a large increase in aboveground competition, with plants in eutrophic conditions needing significantly more aboveground biomass to their requisite aboveground resources.

**Figure 3.**
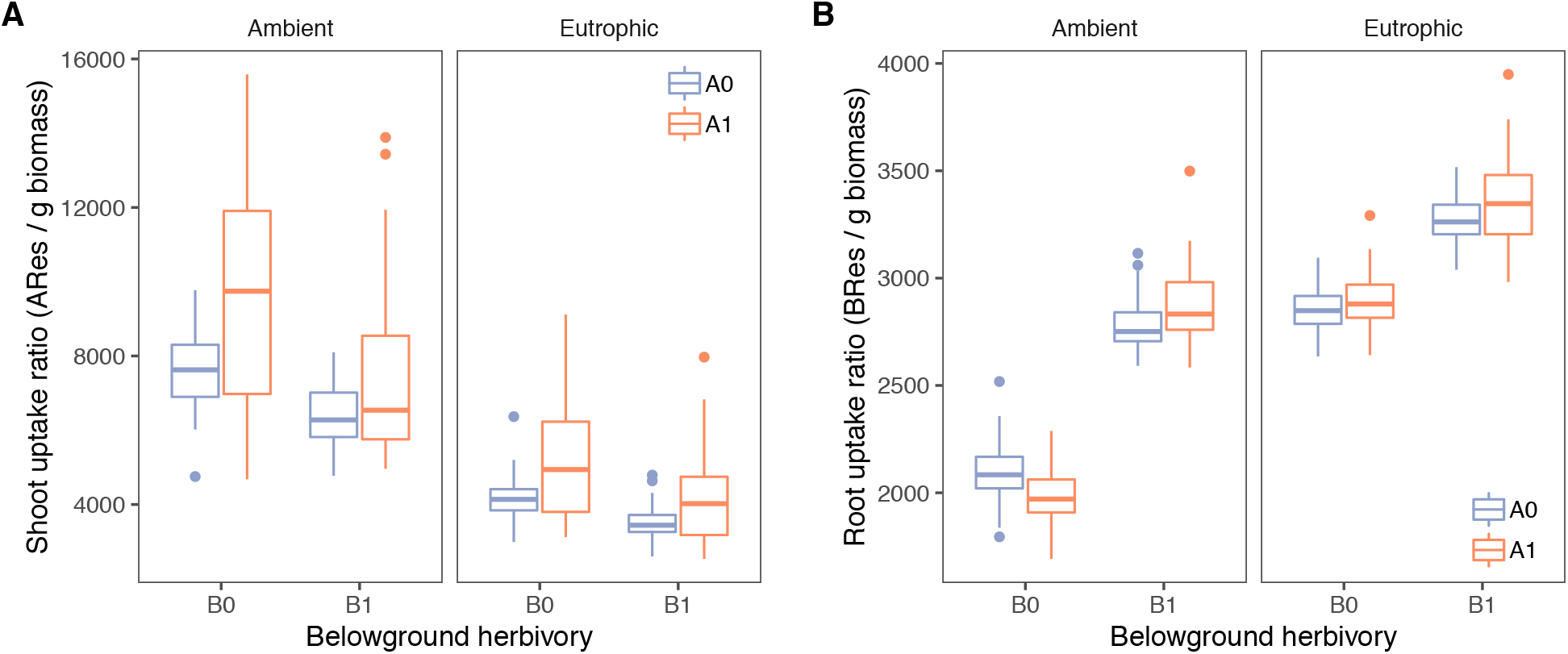
(A) Shoot uptake ratio and (B) root uptake ratio with experimental treatment. Each ratio reflects the resources acquired per unit biomass. An increasing ratio indicates decreasing competition for the respective resource. Boxplots are derived by averaging all plant-individuals’ per replicate, one simulation year before the experimental treatment terminates. A0 – Aboveground herbivory removed, A1 – Aboveground herbivory present; B0 – Belowground herbivory removed, B1 – Belowground herbivory present. While shoot herbivory decreases the amount of competition aboveground, belowground herbivory mimics the influence of eutrophication in shifting competition aboveground.

A complementary pattern emerged when inspecting the average root uptake ratio of each community (𝐵𝑅𝑒𝑠 ∶ 𝑚*_root_*, Fig. 3B). A high root uptake ratio corresponds to a plant taking up many resources per gram root biomass per time step, reflecting low competitive intensity; plants receive much of the resources they can take up. By contrast, a low uptake ratio means that plants only take up few resources for each gram of their root biomass, reflecting an intense competitive environment. Eutrophic conditions dramatically increased the amount of belowground resources taken up per gram biomass, reflecting a decrease in belowground competition. Likewise, introducing belowground herbivores also decreased belowground competition, as removal of root biomass increased the amount of remaining uncontested resources. Introducing aboveground herbivory had a split effect: Without belowground herbivores, it increased the amount of belowground competition, as the reduction in competition aboveground reverberated into the belowground compartment. With belowground herbivory, however, the two effects cancelled out.

### 4.3 Resilience dynamics during eutrophication

We next investigated the two metrics central to resilience: resistance and recovery (Hodgson et al. 2015) with respect to simulated Shannon-diversity. The diagnostic resistance of our model grasslands, i.e. the response of diversity one year after herbivory was removed, was significantly lower when eutrophication was part of the treatment (Fig. 4). Without eutrophication, there was very little immediate change when belowground herbivores were removed, though aboveground herbivore removal was mildly impactful (Table 2). In other words, eutrophication has a very large immediate effect on how herbivory affected diversity, suggesting that herbivory, and its characteristics, are important modulators of diversity in eutrophic grasslands. The only other strongly significant interaction was between eutrophication and the removal of belowground herbivores, with the removal of root herbivores somewhat mitigating the immediate impact of eutrophication.

**Figure 4.**
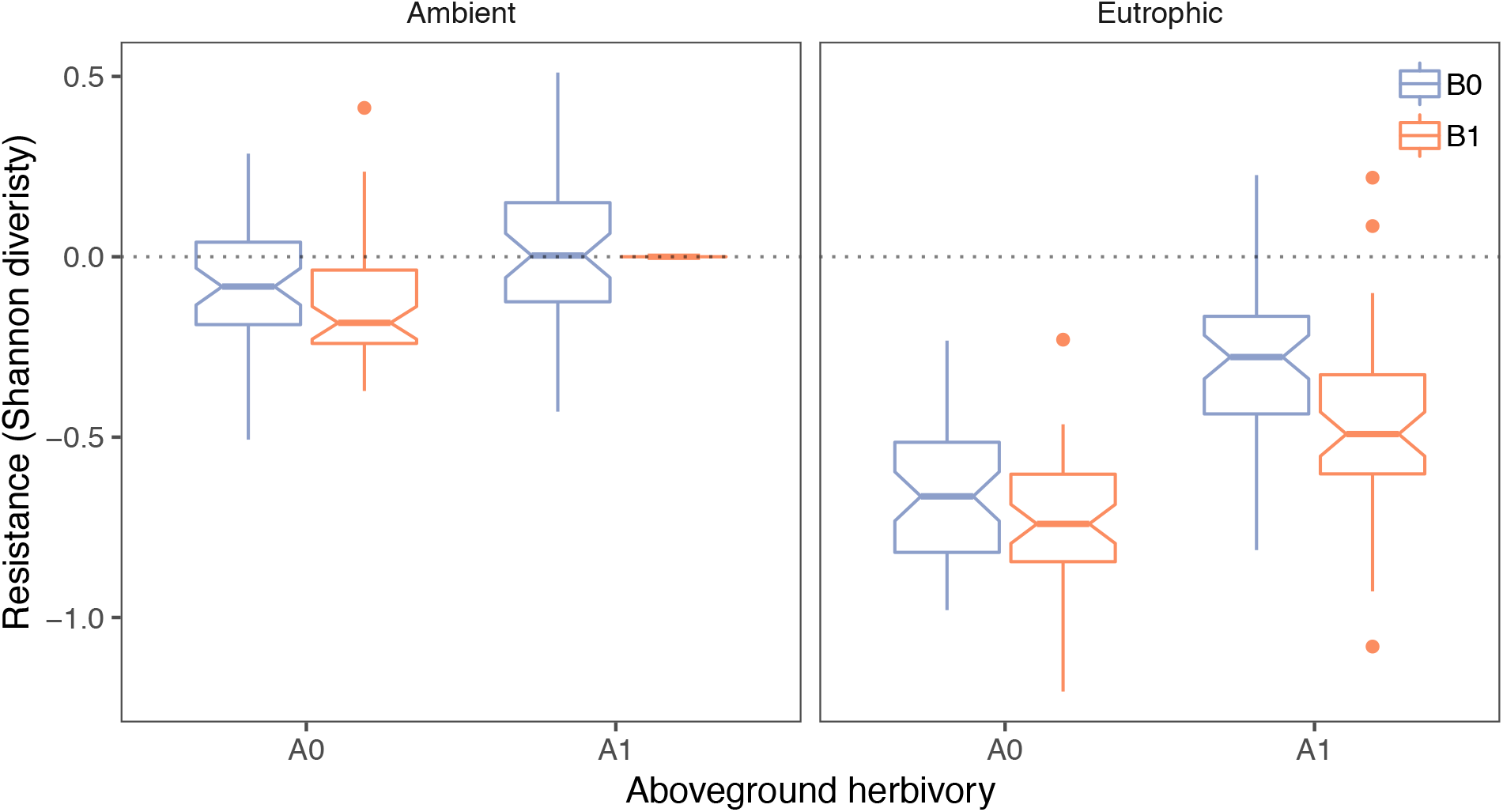
Resistance of Shannon diversity to the experimental treatment. Resistance is measured as the change in Shannon diversity from the control diversity one year after the treatment begins. The control diversity is defined as the treatment maintaining ambient belowground resources and both above- and belowground herbivory. A0 – Aboveground herbivory removed, A1 – Aboveground herbivory present; B0 – Belowground herbivory removed, B1 – Belowground herbivory present. Notches indicate 95% confidence intervals of the medians. While aboveground competition increases the resistance of grasslands to eutrophication, belowground herbivory decreases its effect.

**Table 2:** Impact of above- and belowground herbivory removal on the resistance response of simulated grassland plots to eutrophication.

| Variable | Estimate | Std. Error | t-value | p-value |
| --- | --- | --- | --- | --- |
| Intercept | 1.199e-15 | 0.026 | 0.000 | 1.000 |
| Eutrophication | -0.467 | 0.037 | -12.729 | < **0.001** |
| A0 | -0.131 | 0.037 | -3.578 | < **0.001** |
| B0 | 0.012 | 0.037 | 0.322 | 0.748 |
| Eutrophication:A0 | -0.134 | 0.052 | -2.588 | < **0.05** |
| Eutrophication:B0 | 0.171 | 0.052 | -2.588 | < **0.01** |
| A0:B0 | 0.040 | 0.052 | 0.768 | 0.443 |
| Eutrophication:A0:B0 | -0.149 | 0.073 | -2.035 | < **0.05** |
| Adjusted R-squared: 0.692; F-statistic: 129.1 on 7 and 392 DF |  |  |  |  |

Recovery was measured as the number of years it takes the community to return to its starting Shannon diversity. As expected, the control scenario—with ambient belowground resources and retaining both compartments’ herbivores—was the fastest to recover (Fig. 5), taking on average two years to fall within two standard deviations of its sample mean for ten years (see *Methods: Simulation design and experiments*), in line with a normally distributed test of the algorithm (see Appendix S2: Fig. 3). The removal of belowground herbivores increased the TTR compared to the control, but the most damaging experimental configuration—in terms of TTR—was removing aboveground herbivores yet leaving those belowground. In scenarios with eutrophication, the community is quicker to recover if it had a history of belowground herbivory removal, agnostic of aboveground herbivory.

**Figure 5.**
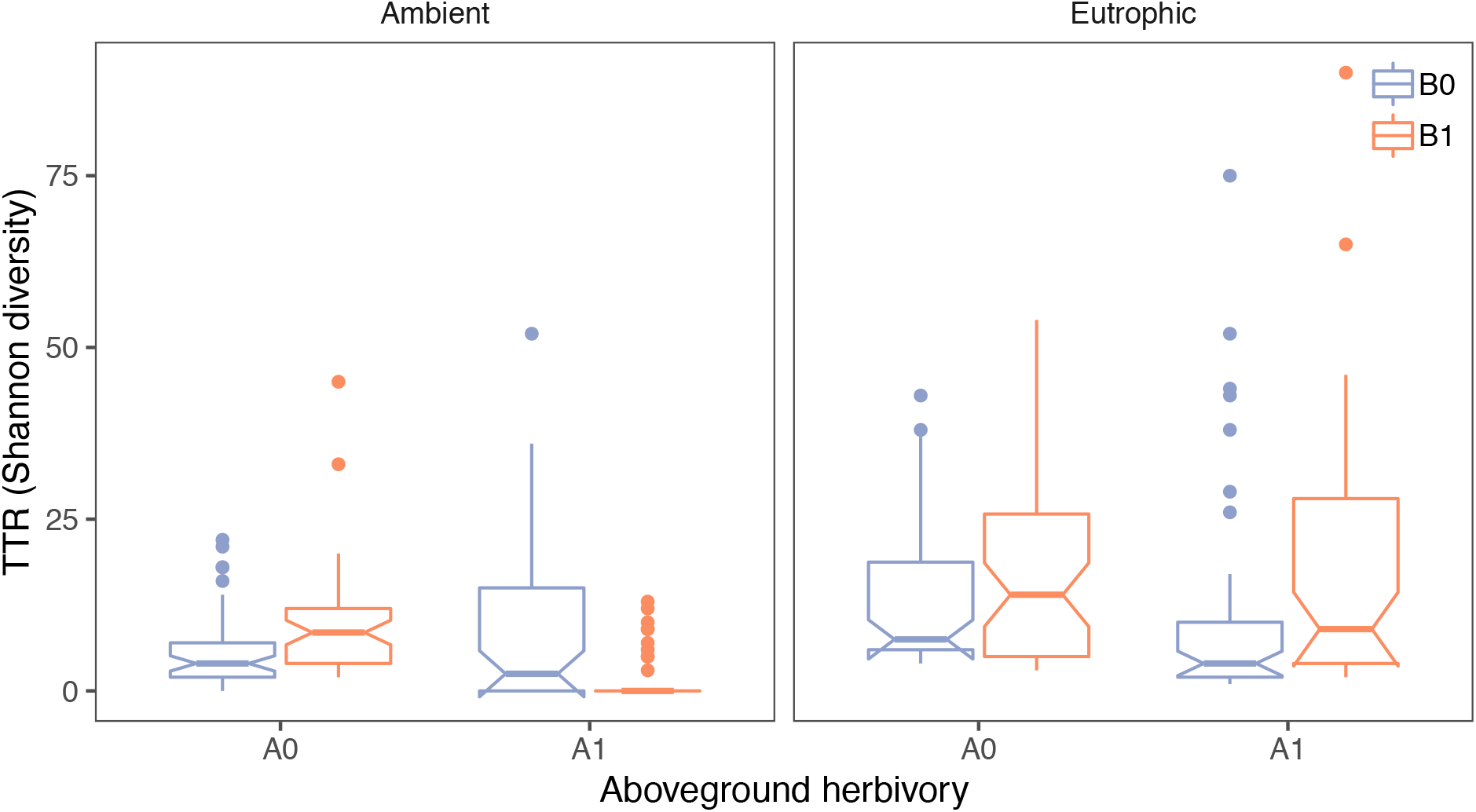
Recovery of Shannon diversity after the experimental treatment. Recovery is measured as the number of years required for Shannon diversity to return to and stay within 2σ of its pre- disturbance level for ten simulation years (time to return, TTR). The control is defined as the treatment maintaining ambient belowground resources and both above- and belowground herbivory. A0 – Aboveground herbivory removed, A1 – Aboveground herbivory present; B0 – Belowground herbivory removed, B1 – Belowground herbivory present. Notches indicate 95% confidence intervals of the medians.

Analyzing the impact of the belowground herbivore’s preferentiality (𝛼) on the TTR of Shannon diversity, we found that there is little difference between the different herbivory regimes aside from one effect: with ambient belowground resources, the presence of generalist herbivores slightly reduced the time to return compared to the various levels of preferential herbivory, though its median stayed the same (Appendix S2: Fig. 4).

## 5 Discussion

Although the interaction between grassland eutrophication and aboveground herbivory has driven significant interest (Borer et al. 2014, Anderson et al. 2018), the relationship between eutrophication and belowground herbivory has remained largely unexplored. Eutrophication tends to decrease species richness by shifting competition from the belowground compartment to the aboveground compartment, giving a disproportionate advantage to fast growing, tall, species (Bobbink et al. 1998; Hautier 2009; Farrer et al. 2016).

In our analysis, using a simulation model we factorially removed above- and belowground herbivores and introduced eutrophication. We found that belowground herbivory compounds the impact of eutrophication on diversity, because like eutrophication it increases the relative proportion of belowground resources to root biomass, resulting in more resources available to the plants’ remaining roots. This shifts competition to the aboveground compartment, exacerbating the advantage of largest, fastest growing plants.

The clearest evidence of this dynamic within IBC-grass is the relative change in the plants’ uptake ratios (Fig. 3). Disregarding above- and belowground herbivory, eutrophication *per se* increases the amount of belowground resources available to the plants, increasing the resources available per gram of remaining root biomass. This new resource abundance shifts competition aboveground, leading to the competitive exclusion of short and slow-growing species. These dynamics are consistent with the most contemporary understandings of eutrophication in grasslands (DeMalach et al. 2016, 2017). As these dynamics were not imposed within the model’s design (May et al. 2009; Weiss et al. 2014), they can be considered an independent, successful prediction—a strong indicator of structural realism of IBC-grass, defined as its potential to realistically capture the key elements of a grassland’s internal organization (Wiegand et al. 2003; Grimm and Berger 2016).

Our analysis of belowground herbivory suggests that its main effect is to also increase the amount of resources available to the plants’ roots; rather than increasing the amount of nutrients in the soil, root herbivores decrease the amount of biomass present to compete for it. This synthesis of the theory behind eutrophication and the potential impacts of belowground herbivores is new, and has very little footprint within the literature, with only two empirical studies previously examining it.

In the first, La Pierre et al. (2014), found that removing belowground herbivores increased plant species evenness after eutrophication events, finding that they behaved as a top-down control on the sub-dominant species within the grassland. Once the herbivores were removed, these species were also able to utilize the new resources and diversity increased. In the second, the relationships between above- and belowground herbivory and eutrophication were also examined empirically by Borgström et al. (2017)—the template for our study’s design. They found that both belowground herbivores and eutrophication depress diversity, while aboveground herbivory increases it. Although these general effects are consistent across both our studies, the smaller interactions are not fully consistent. While Borgström et al. (2017) also found that root herbivory generally decreased species evenness, they found that the herbivores’ effects were more pronounced with ambient resource levels. As these two experiments represent very different study systems, some discrepancies between them are not surprising. As a simulation model, IBC-grass enables us to observe the diversity patterns of a simplified grassland, emergent from basic ecological assumptions. Since Borgström and colleagues used grassland mesocosms, their results will incorporate nuances endemic to their grassland system. This degree of detail, however, may obscure the larger picture; while models’ simplifications inevitably omit some of the more complex interactions embedded in real grasslands, simplification enables us to examine the processes likely general to most grassland ecosystems.

Several secondary factors confound the direct comparison of the two studies. IBC-grass uses a larger spatial extent and much larger species pool, and its aboveground herbivory is modeled after ungulates rather than invertebrates. Further, while Borgström et al. (2017) could precisely measure the amount of nitrogen added to the soil, in IBC-grass belowground resources are phenomenological and correspond simply to the “resources taken up belowground,” and therefore could be water or nutrients. Therefore, it is not surprising that these systems react somewhat differently to our experimental treatments. However, the main effects found in the empirical study could be replicated without fine-tuning the model’s parameters: Belowground herbivory and eutrophication generally negatively impact diversity while aboveground herbivory increases it.

### 5.1 Root herbivory’s influence on symmetric competition for belowground resources

Setting aside eutrophication *per se*, given root herbivores’ prevalence in grasslands (Kozlov and Zvereva 2017), building a theoretical understanding of their impact is critical to understanding these systems’ diversity. There is little consensus on how herbivory belowground could change a grassland’s diversity, with studies finding its influence anywhere from negative (Brown and Gange 1989A, 1989B; Fraser and Grime 2001; Körner et al. 2014) to positive (De Deyn 2003; Stein et al. 2010; Borgström et al. 2017), to neutral (Wurst and Rillig 2011). We argue that this variation in root herbivory’s effect stems from two processes: The preferentiality of the root herbivores and the (a)symmetry of belowground competition itself. Our results suggest that for root herbivores to positively impact grassland diversity, belowground competition must be asymmetric. When it is not, even very preferential herbivores will likely only neutrally impact diversity.

For a resource like light, herbivores reduce the competitive asymmetry between the largest and smallest plants by generating space in the upper canopy, feeding the plants lower to the ground (Borer 2014; Anderson 2018). However, as belowground competition is often symmetric, our results suggest that herbivores will reduce the plants’ biomasses without equalizing their competitive fitness. Any decrease in root biomass will only result in an excess of belowground resources per remaining root biomass, thus reducing root competition. As the belowground compartment is no longer limiting, aboveground competition will increase, and because aboveground competition is asymmetric, its increase will consequently lead to the exclusion of shorter, slower growing species.

To contextualize this result, it helps to compare the mechanisms behind belowground herbivory to those aboveground. Compared to root herbivory, the impact of shoot herbivory is composed of two asymmetries: The largest plants are eaten asymmetrically, but crucially this reduction in the plants’ sizes minimizes a competitive asymmetry between large and short individuals. With symmetric belowground competition, by contrast, no such competitive asymmetry exists. Therefore, root herbivores will generally not be as effective in maintaining biodiversity as their aboveground cousins.

If belowground competition is symmetric, our results further indicated that even extremely preferential root herbivory may be unable to increase diversity compared to scenarios without root herbivores. This reflects a balance between the positive influence of disproportionately reducing the largest plants’ root biomasses and the corresponding increase in aboveground competition. Of course, relative to purely generalist herbivores, even a low degree of feeding preference will increase diversity (Fig. 2). This suggests that if belowground competition happens to asymmetric, it is likely that any preference of the root herbivores towards dominant plant functional types could prove to be a strong mechanism maintaining a grassland’s diversity. Given that other empirical analyses of root herbivory suggest that it may stabilize diversity (De Deyn 2003; Stein et al. 2010; Borgström et al. 2017), future research should investigate the possibility of underlying asymmetries in belowground competition within these study systems.

That belowground resources are symmetrically allocated has been an important assumption in grassland and forest modeling (Schwinning and Weiner 1998). Although numerous empirical tests have found this to be the case (Casper and Jackson 1997; Weiner 1997; Berntson and Wayne 2000; Blair 2001; Wettberg and Weiner 2003; Cahill and Casper 2003; Lamb 2008), under certain conditions it is likely to be asymmetric as well (Weiner 1990; Rajaniemi 2002, 2003; Rajaniemi et al. 2003, Weiss et al. 2019). Understanding when belowground herbivory is likely to be asymmetric is therefore necessary to fitting our model’s results into a broader context.

The empirical literature has found that asymmetry in belowground competition is increased when nutrients are patchy in the soil, giving a competitive advantage to larger root systems that can disproportionately access them (Fransen et al. 2001; Facelli and Facelli 2003; Rajaniemi 2002; Rajaniemi et al. 2003). A model of belowground competition (Raynaud and Leadley 2005) furthered this hypothesis, finding that the symmetry of belowground competition should also depend on how nutrients diffuse in the soil. In wet soils, nutrients will be more diffuse and therefore plant competition will become more dependent on the plants’ uptake kinetics, shifting competition towards asymmetry. This hypothesis has been supported by at least one empirical test (Rewald and Leuschner 2009). Belowground competition could also become asymmetric through its vertical dimension (Schenk 2006), with root systems higher to the surface receiving a larger proportion of water and nutrients (van Wijk and Bouten 2001).

To summarize, although belowground competition is often more symmetric than aboveground competition, this balance should not be taken for granted. When belowground competition is symmetric, our model indicates that root herbivory will not support diversity and may even substantially decrease it as it shifts competition aboveground, leading less competitive species towards exclusion. Given that even an extreme asymmetry in the feeding preferences of the herbivores did not shift the direction of their influence on coexistence, our results indicate that variation in the empirical literature on root herbivores likely results not from their feeding preferences in isolation, but also from asymmetries in belowground competition.

### 5.2 Resilience of grassland systems to eutrophication

Trophic interactions, such as herbivory, are acknowledged as important contributors to the stability of ecological systems through their compensatory effects (Hillebrand et al. 2007; Gruner et al. 2008; Ghedini et al. 2015). To supplement our investigation into the mechanisms through which herbivory influences grasslands’ responses to eutrophication, we also measured how above- and belowground herbivory change the resistance and recovery of our model grasslands to eutrophication. Our main findings indicate that the removal of herbivores is relatively mild in its immediate effect on diversity (Fig. 4), and that once the herbivores return diversity follows relatively quickly (Fig. 5).

With eutrophication, however, the magnitude of change is much larger. For resistance, as predicted by ecological theory (Hillebrand et al. 2007; Gruner et al. 2008; Kohli et al. 2019) and empirical evidence (Eisenhauer et al. 2011; Post 2013; Ghedini et al. 2015; McSkimming et al. 2015), the presence of aboveground herbivory compensates—albeit weakly—for the immediate effects of a strong eutrophication event (Fig. 4), and over the course of the 100-year treatment, its impact becomes much more influential (Fig. 1). This suggests that a long experimental duration is necessary to understand the full array of interactions forming a grassland’s response to eutrophication (Kidd et al. 2017). Further, simulations retaining aboveground herbivores returned to their pre-disturbance state more quickly than those without it (Fig. 5).

Belowground herbivory, however, does not neatly dovetail with ecological theory: Indeed, the presence of root herbivory coincides with a reduction in resistance to eutrophication (Fig. 4), as well as a longer time to return (Fig. 5). This is unsurprising, however, given our finding that belowground herbivores tend to exacerbate the dominance of the strongest competitors, unlike their counterparts aboveground. Once established, these plant species will retain their dominance for a long period of time after nutrient addition is halted.

## 6 Conclusions

As anthropogenic changes such as eutrophication increasingly stress grassland ecosystems, understanding what aspects of these communities mediate their ability to resist degradation is becoming increasingly important. Trophic interactions between the plant community and their herbivores are one such aspect. Our results suggest that rather than strengthening a plant community’s resilience to a eutrophication event, belowground herbivores compound its negative influence on plant diversity and resilience. These results are tightly interlinked with the symmetry of belowground competition and preferences of the herbivores themselves. Future research must investigate how prevalent competitive asymmetries are within the belowground compartment, as they may be a necessary condition for root herbivores to positively influence diversity.

## Supporting information

AppendixS2

AppendixS1

## 7 Author contributions

FM developed, with FJ and VG, the original model; MC and FJ formulated the research question; MC performed, with UES and FJ, the analysis and was the lead author; IS and SW added expert knowledge on belowground herbivory; all authors contributed to the interpretation of results and the writing of the manuscript.

## 8 Acknowledgements

MC, with FJ, were funded by DFG Priority Program 1374, “Infrastructure-Biodiversity-Exploratories” (DFG-JE 207/5-1). UES was supported by Deutsche Forschungsgemeinschaft in the framework of the BioMove Research Training Group (DFG-GRK 2118/1). SW was funded by the DFG Priority Program 1374, “Infrastructure-Biodiversity-Exploratories” (DFG-WU 603/3-3).

We thank the managers of the three Biodiversity Exploratories and all former managers for their work in maintaining the plot and project infrastructure, Christiane Fischer, Anja Hoeck, and Cornelia Weist for giving support through the central office, and Markus Fischer, Eduard Linsenmair, Dominik Hessenmöller, Daniel Prati, Ingo Schöning, François Buscot, Ernst-Detlef Schulze, Wolfgang W. Weisser and the late Elisabeth Kalkofor for their role in setting up the Biodiversity Exploratories project.

We lastly thank Niv DeMalach for helpful conversations about the interaction of competition and eutrophication that helped to develop this work.

