## AppendixS2 for "While shoot herbivory mitigates, root herbivory exacerbates eutrophication’s impact on diversity in a grassland model"

**Supplement B: Additional graphs**

### 1 Replication of Borgström et al. (2017)—Pielou's Evenness (E)

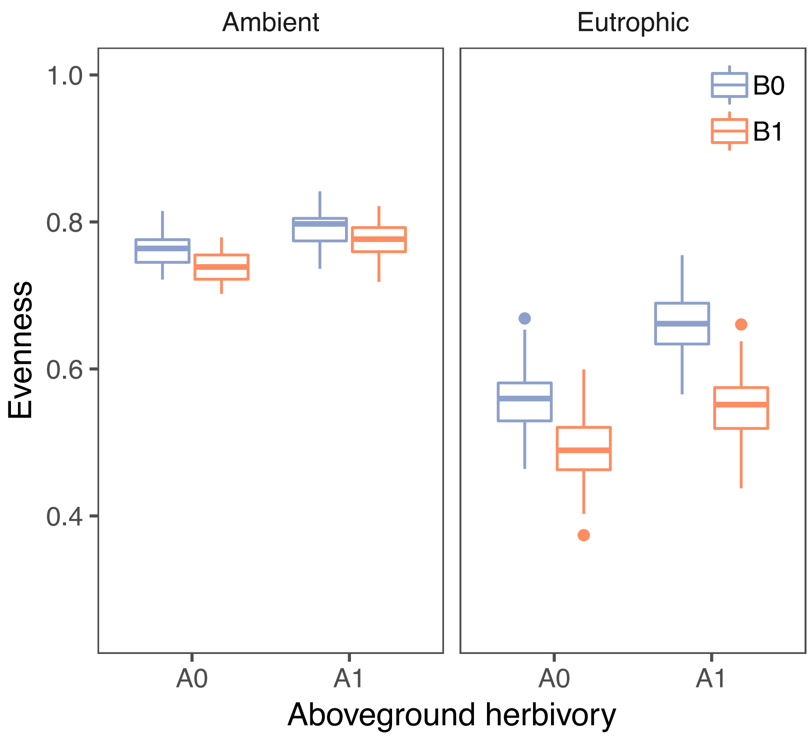


Figure B1: Pielou's evenness after three years of experimental treatment. Borgstrom et al. (2017) calculates Pielou's evenness, rather than Shannon diversity, in their mesocosm experiments. This variable differs from raw Shannon diversity in that it is normalized to log(S), where S is the number of species present in the community. Although numerous differences between study systems will complicate the direct comparison of our data to theirs, the main interactions remain consistent: Aboveground herbivores help maintain diversity, while belowground herbivores and eutrophication generally diminish it. One of the central differences between these two datasets is the dramatically higher evenness seen in the ambient resource simulations. The default resource setting of IBC-grass (60 BRes, 100 ARes) generally leads to rich (~ 40 species/$m^2$) communities. For ambient resources, while above- and belowground herbivory treatments alter the number of species within the plots, they do not influence the evenness of the resultant communities nearly as much. A0 – Aboveground herbivory removed, A1 – Aboveground herbivory present; B0 – Belowground herbivory removed, B1 – Belowground herbivory present.

*Borgström, P., J. Strengbom, L. Marini, M. Viketoft, and R. Bommarco. 2017. Above- and belowground insect herbivory modifies the response of a grassland plant community to nitrogen eutrophication. Ecology 98:545–554.*

### 2 Effect of herbivore preference on the dominant PFT’s total root biomass

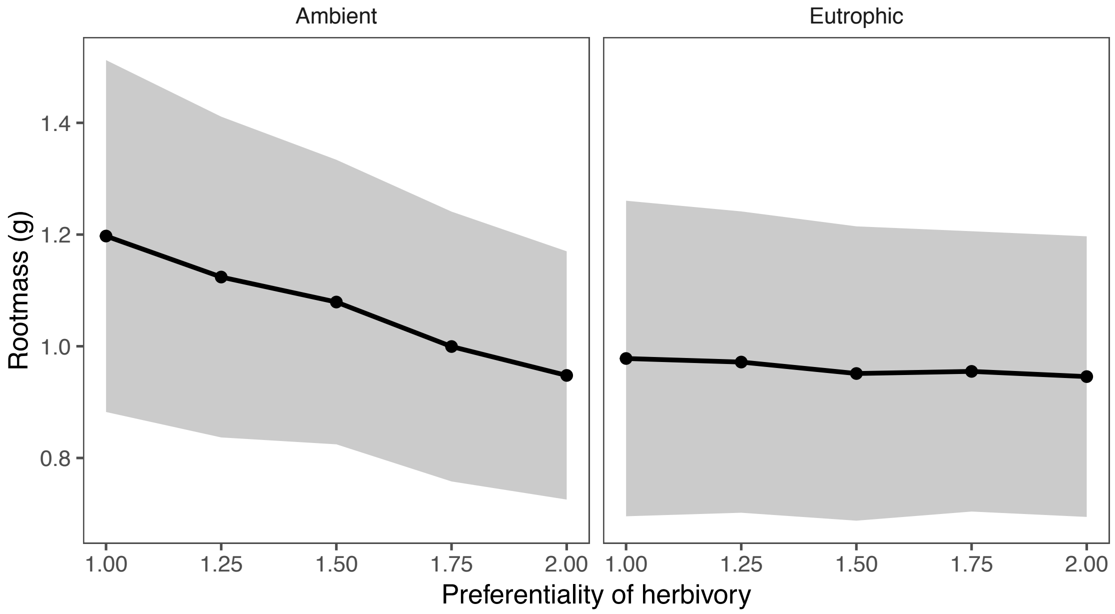


Figure B2: Change in root biomass of the dominant PFT, defined as the PFT with the most root biomass in simulations with generalist belowground herbivores. Ribbons indicate 1σ around the mean.

### 3 Standardized test of recovery algorithm

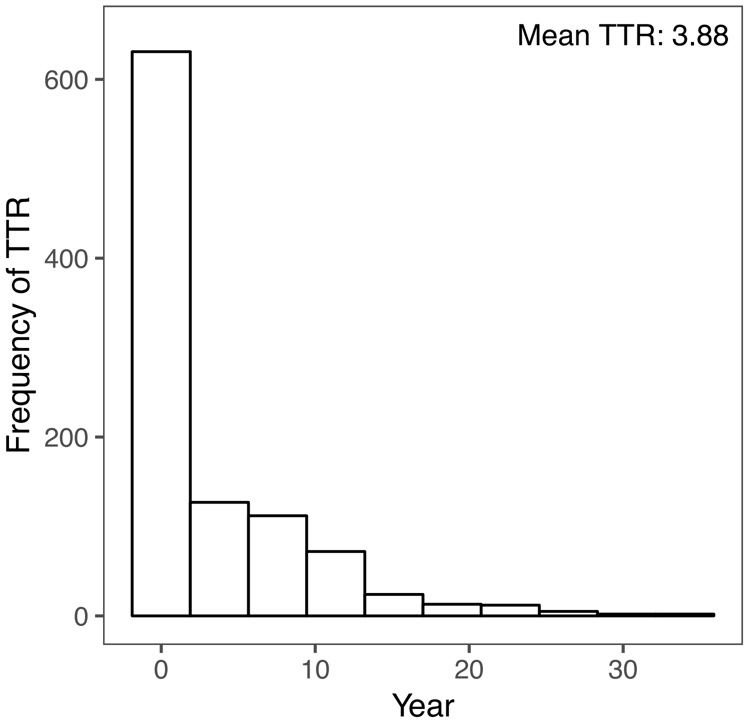


Figure B3: Results of running the recovery algorithm (deriving the TTR) on a timeseries dataset of 1000 replicates derived from sampling a Gaussian distribution ($\mu = 0$, $\sigma = 1$). The average number of years required for a replicate to return to within $2\sigma$ of the mean and stay within this window for 10 years was 3.8 years.

### 4 Effect of herbivore preference on recovery

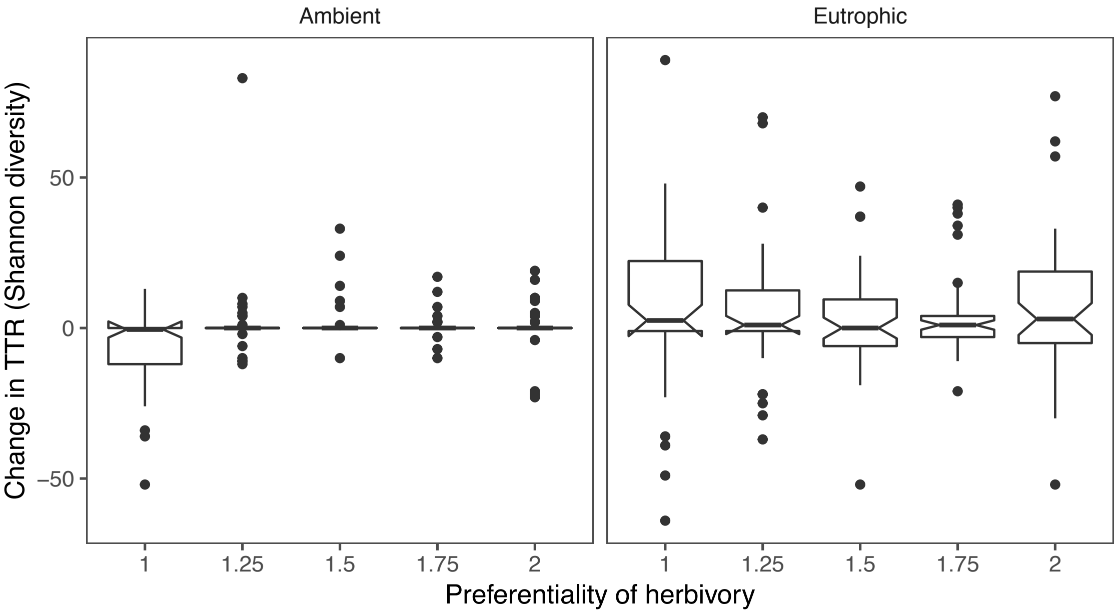


Figure B4: Change in time to return (measured in years; TTR) in Shannon diversity between simulations with belowground herbivores and those without. Aboveground herbivory is set to present; Notches reflect 95% confidence intervals of the median.
