## AppendixS1 for "While shoot herbivory mitigates, root herbivory exacerbates eutrophication’s impact on diversity in a grassland model"

### Supplement A: The individual-based community grassland model—IBC-grass

#### Detailed model description

The model description follows the ODD protocol (Overview, Design concepts, Detail) for describing individual-based models (Grimm et al. 2006, Grimm et al. 2010). The description of the original model by May et al. (2009) is coloured black. Modifications by us are marked red.

#### 1 Purpose

The model is designed to evaluate the response of plant functional type diversity towards aboveground grazing and belowground grazing, during and after eutrophication disturbances.

#### 2 Entities, state variables and scales

The model includes the entities seeds, individual plants, and grid cells (Table A1). Seeds are described by the state variables position, age, and mass. Plant individuals are characterized by their position, the mass of three plant compartments (shoot, root, and reproductive mass), the duration of resource stress exposure, and they are classified as a certain PFT with specific trait attribute parameters (see May et al. 2009 for the parameterization for the PFTs). Spatially, plants are described by their "zone-of-influence" (ZOI), i.e. a circular area around their location (Schwinning and Weiner 1998, Weiner et al. 2001). Within this area the individual can acquire resources and if the ZOIs of neighboring plants overlap, the individuals will only compete for resources in the overlapping area. For the three-compartment model version we consider two independent ZOIs for a plant's shoot and root, representing above- and below-ground resource uptake and competition. The ZOIs radii are determined from the biomass of the corresponding plant compartment.

In order to simplify spatial calculations of resource competition, ZOIs are projected onto a grid of discrete cells. Grid cells represent 1 $cm^2$. The state of a grid cell is defined by two resource availabilities, above and below ground. The size of the modelled area was 128 cm × 128 cm. To avoid edge effects, periodic boundary conditions were used, i.e. the grid essentially was a torus. Through periodic boundaries, we ensure that dispersal limitation is the same for individuals in the center and at the edge of the simulated area. A model's time step corresponds to one week; a vegetation period consisted of 30 weeks per year, and simulations were run for 100 years.

### 3 Process overview and scheduling

The processes resource competition, plant growth and plant mortality are considered within each week of the vegetation period. Seed dispersal and seedling establishment are limited to certain weeks of the year (Table A2). Aboveground herbivory—grazing—events occur randomly with a fixed probability which is constant for all weeks while belowground herbivory occurs on a weekly schedule. Eutrophication occurs during the middle of the experimental period, from simulation years 100-200. Two processes, winter dieback of above-ground biomass and mortality of seeds are only considered once a year, at the end of the vegetation period (Fig. A2). Plant's state variables are synchronously updated within the subroutines for growth, mortality, grazing and winter dieback, i.e. changes to state variables are updated only after all model entities have been processed (Grimm and Railsback 2005).

### 4 Design concepts

#### 4.1 Basic concepts

The model represents trait-offs in the plants' traits, and it focusses on plant function types (PFT), not on predefined species. Community assembly is mimicked by starting with all possible combinations of trait combinations and letting the final community composition assemble itself. Different modes of competition, i.e. asymmetric and symmetric, are implemented for above- and below-ground competition, respectively. Adaptive resource allocation is implemented (see below, Design concept "Adaptation".

#### 4.2 Emergence

All features observed at the community level, such as community composition and diversity, emerged from individual plant-plant interactions, grazing effects at the individual scale, and resource availabilities.

#### 4.3 Adaptation

In the submodel representing plant growth and above- and below-ground competition, plants adaptively allocate resources to shoot and root growth in order to balance the uptake of above- and below-ground resources (see below, Submodels – Plant growth and mortality).

#### 4.4 Interaction

Competitive interactions between plant individuals were described using the ZOI approach.

#### 4.5 Stochasticity

Seed dispersal and establishment, as well as mortality of seeds and plants are modelled stochastically to include demographic noise. Grazing events occur randomly during the vegetation period and the affected plants are chosen randomly, but the individual's probability of being grazed depends on plant traits (see Submodels – Grazing).

#### 4.6 Observation

To describe community diversity and composition the individual numbers of all PFTs are recorded each year in week 20 directly before seed germination in autumn. These data are used to calculate annual values of Shannon diversity, which contains more information about the distribution of relative abundances among PFTs than the number of surviving PFTs only. Additionally, the overall above- and belowground biomasses are recorded to distinguish the responses of type richness and standing biomass.

### 5 Initialization

Initially, ten seedlings of all 81 PFTs (see section Plant traits and PFT parameterization below) with their respective seedling mass were randomly distributed over the grid. Their germination probability was set to 1.0 to assure equal initial population sizes of all PFTs. A spatially and temporally homogenous distribution of resources (both above- and below-ground) was used in all simulation experiments.

### 6 Input data

The model does not include any external input of driving environmental variables.

### 7 Submodels

#### 7.1 Competition

Following the ZOI approach, plants compete for resources in a circular area around their central location point. To relate plant mass to the area covered ($A_{shoot}$), we extended the allometric relation used by (Weiner et al. 2001)

$$A_{shoot} = SLA \cdot (LMR \cdot m_{shoot})^{2/3} \qquad \text{(A1)}$$

where SLA is a constant ratio between leaf mass and ZOI area and $m_{shoot}$ is vegetative shoot mass (compare Tables A1 and A2). The LMR is introduced to describe different shoot geometries and is defined as the proportion of photosynthetically active (leaf) tissue to the total (shoot) tissue (Fig. A1). The allometric relationship between the amount of photosynthetic biomass and the area of the ZOI is consistent with the model of Weiner et al. (2001), eq. 1, and presents sigmoidal growth in the absence of competition. Given the phenomenological nature of this relationship, we follow Weiner et al. 2001 and omit unit-rectifying constants from this and each subsequent equation within the model. Only the former is considered for the calculation of the ZOI size. These circular areas are projected onto a grid of discrete cells. Grid cells thus contain the information by which plants they are covered, so that resource competition can be calculated cell by cell. The resources within a cell are shared among plants according to their relative competitive effects ($\beta_i$). The resource uptake ($\Delta$res) of plant i from a cell with resource availability ($Res_{cell}$) covered by n plants is thus calculated as

$$\Delta res_i = \frac{\beta_i}{\sum_{j=1}^{n} \beta_j} \cdot Res_{cell}. \qquad \text{(A2)}$$

Calculating $\beta_i$ in different ways allows including different modes of competition (Weiner et al. 2001). We assume that the relative competitive ability of a plant is correlated with its maximum growth rate in the absence of resource competition. Therefore, $\beta_i$ is proportional to maximum resource utilization per unit area covered ($g_{max}$), see Submodel: Plant growth and mortality and Table A2), In the case of size-symmetric competition, $\beta_i$ simply equals $g_{max}$:

$$\beta_i = g_{max} \qquad \text{(A3a)}$$

In the case of partially size-asymmetric competition $\beta_i$ is a function of plant mass and shoot geometry:

$$\beta_i = g_{max} \cdot m_{shoot} \cdot LMR^{-1} \quad \text{(A3b)}$$

The inverse of LMR is used, because plants with a lower fraction of leaf tissue are considered to be higher and thus show a higher competitive ability by overtopping other plants (Fig. A1). In this way, plants with equal $g_{max}$ receive equal amounts of resources from one unit of area irrespective of their mass or height in the case of size-symmetric competition, while larger and higher plants receive a higher share of resources in proportion to their shoot geometry traits in the case of partially asymmetric competition (Schwinning and Weiner 1998, Weiner et al. 2001). The resource uptake of one plant within one week can then be determined by summing the results of Eq. (A2) over all cells covered by the plant.

To include differences between intra- and interspecific competition, individuals of the same PFT are considered as conspecifics and those of different PFTs as heterospecifics. The relative competitive ability βi of one plant is then determined as a decreasing function of the number of plants belonging to the same PFT ($n_{PFT}$) and covering the same cell:

$$\beta_i = g_{max} \cdot \frac{1}{\sqrt{n_{PFT}}} \quad \text{(A3c)}$$

Eq. (A3c) is used for size-symmetric competition instead of Eq. (A3a). In the case of size asymmetry, plant mass and geometry are taken into consideration according to Eq. (A3b). This approach represents a situation where intraspecific competition is increased relatively to interspecific competition and therefore implicitly includes niche differentiation of resource competition at the cell scale, which has been known as an important factor for species coexistence (Chesson 2000, Silvertown 2004). In the model analysis, versions with and without niche differentiation were compared in order to test if this assumption for competition at the cell scale translates into a different behavior at the community scale (results see May et al. 2009).

#### 7.2 Plant growth and mortality

Plant growth only depends on the resources (Δres) that the plant acquired during the current time step. In the absence of competition, plants show sigmoidal growth (Hunt 1982). Therefore, we again adapted the allometric growth equation used by Weiner et al. (2001) to the description of plant geometry used here:

$$\Delta m = g \cdot \left( \Delta res - SLA \cdot LMR^{\frac{2}{3}} \cdot g_{max} \cdot \frac{m_{shoot}^{2}}{m_{max}^{\frac{4}{3}}} \right) \quad \text{(A4)}$$

where g is a constant conversion rate between resource units and plant biomass and $m_{max}$ is the maximum mass of shoot or root, respectively. In addition, the maximum amount of resources that is allocated to growth each week is limited by a maximum resource utilization rate given by $g_{max}$ [resource units/cm²] multiplied by ZOI area [cm²]. If Eq. (A4) yields a negative result, Δm is set to zero and thus negative growth is prohibited. The terms were designed in Weiner et al. (2001) such that: 1) growth is proportional to the resources available to the plant, scaling with the ZOI area, 2) without competition, growth is sigmoidal, 3) the equation is as simple as possible.

Growth of reproductive mass is restricted to the time between weeks 16 – 21. In this period, a constant fraction of the resources (5% for all PFTs) is allocated to growth of reproductive mass (Schippers et al. 2001, Kahmen 2004), and reproductive mass is limited to 5% of shoot mass in total. The same resource conversion rate (g) is used for reproductive and vegetative biomass.

Eqs. (A1) – (A4) are applied to shoot and root ZOIs independently, with the difference that for root growth the factor LMR is always one. We assume that the minimum uptake of above- and below-ground resources limits plant growth (Lehsten and Kleyer 2007) and introduced adaptive shoot-root allocation in a way that more resources are allocated to the growth of the plant compartment that harvests the limiting resource (Shipley and Meziane 2002, Weiner 2004). For resource partitioning, we adopt the model of Johnson (1985) and the fraction of resources allocated to shoot growth is calculated as

$$\alpha_{shoot} = \frac{\Delta res_B}{\Delta res_B + \Delta res_A} \quad \text{(A5)}$$

where $\Delta res_A$ is above-ground and $\Delta res_B$ is below-ground resource uptake. This transport-resistance partitioning model was taken from Johnson 1985 (eq. 16) which relates the root:shoot ratio to the C:N ratio in the plants. Eq. A5 ensures that the ratio of total resources taken up (both above- and belowground) going to the shoots is equal to the rate of resources taken from the soil. For the roots, the proportion is vice versa. This keeps the growth of both compartments balanced.

Plants suffer resource stress if their resource uptake (in any layer) is below a fixed threshold fraction ($thr_{mort}$) of their optimal uptake, which is calculated as maximum resource utilization times ZOI area. That means each week the condition $\Delta res < thr_{mort} \cdot A_{\frac{shoot}{root}} \cdot g_{max}$ is evaluated and if it is true either for shoot or root the plant is considered as stress exposed during this week. Consecutive weeks of resource stress exposure ($w_{stress}$) linearly increase the probability of death

$$p_{mort} = p_{base} + \frac{w_{stress}}{surv_{max}} \quad \text{(A6)}$$

where $surv_{max}$ is the maximum number of weeks a plant can survive under stress exposure and $p_{base}$ is the stress independent background mortality of 0.7% per week corresponding to an annual mortality rate of 20% (Schippers et al. 2001). It's possible for $p_{mort}$ to exceed 1, but the plant will be killed that time step.

Dead plants do not grow and reproduce anymore, but they still can shade others and are therefore still considered for competition in the one-layer model and for at least above-ground competition in the two-layer model. Each week the mass of all dead plants is reduced by 50% and they are removed from the grid completely as soon as their total mass decreases below 10 mg.

#### 7.3 Aboveground herbivory

Grazing is modelled as partial removal of an individual's above-ground biomass. The frequency of grazing is specified by a constant weekly probability ($p_{graz}$) of a grazing event. Grazing is a process that acts selectively towards trait attributes such as shoot size and tissue properties. Therefore, for each plant the susceptibility to grazing ($s_{graz}$) is calculated as a function of shoot size, geometry and PFT-specific palatability (palat).

$$s_{graz} = m_{shoot} \cdot LMR^{-1} \cdot palat \quad \text{(A7)}$$

The probability for each plant to be grazed within one week is derived by dividing individual susceptibilities by the current maximum individual susceptibility of all plants (in other words, the susceptibility of the most-susceptible plant). All plants are checked for grazing in random order. In case a plant is grazed, 50% of its shoot mass and its complete reproductive mass are removed. The random choice of plants is repeated without replacement, until 50% of the total (above-ground) biomass on the whole grid has been removed. When all plants have been checked for grazing once, but less than 50% of the total above-ground biomass has been removed, grazing probabilities for all individuals are calculated once more based on Eq. (A7) and the whole procedure is repeated until 50% of above-ground biomass has been removed or until a residual biomass is reached which is considered ungrazable. This fraction is set to 15 g/m² following (Schwinning and Parsons 1999). This allows a plant individual to be grazed never or several times during one week with a grazing event.

In addition to stochastic grazing, each year at the end of the vegetation period 50% of the above-ground mass of all plant individuals is removed to mimic vegetation dieback in winter.

### 7.4 Belowground herbivory

Belowground herbivory was implemented such that each time step some percentage of the extant biomass is removed from each of the plants, with a gradient of preference in root size ranging from generalist to preferential (i.e. disproportionally eating larger root systems). This herbivory algorithm is intended to reflect the influence of belowground, invertebrate herbivores, such as those belonging to the genus *Agriotes*, one of the most abundant root herbivores in Europe. As this genus generally tends to eat plants with high biomass and growth rates (Sonnemann et al. 2012, 2015), we refrain from explicitly modelling the plants' roots palatability.

The feeding need at week $t$, $n_t$, is calculated as a defined percent (feeding rate, $f$) of that week's expected root mass, which is estimated by averaging each previous week's total realized root biomass $R_i$ for the previous $w$ weeks,

$$n_t = f \cdot \frac{\sum_{i=t-w}^{t-1} R_i}{w} \quad \text{(A8)}$$

For this analysis, the feeding rate ($f$) is 0.1 per week, potentially lower than the typical belowground herbivory pressure (Zvereva and Kozlov 2012), but equal to the aboveground herbivory pressure commonly used in IBC-grass. The number of weeks used to estimate the expected root mass, $w$, is 10. Both parameters are held constant in the following analysis.

The biomass to be removed from each individual's root mass ($g_{i,t}$) is calculated each week as:

$$g_{i,t} = \left(\frac{r_{i,t}}{R_t}\right)^{\alpha} \cdot n_t \quad \text{(A9)}$$

where $r_{i,t}$ is the expected root mass of individual $i$ in week $t$ and $R_t$ is the week's realized total root mass, which may differ from the expected root mass (Fig. A3). $R_t$ differs from $R_i$ in eq. A8 in that $R_i$ refers to the total realized root biomass on week $i$ (and ranging backwards by $w$ weeks), whereas $R_t$ refers to the total realized root biomass on the current week. The parameter $\alpha$ represents the generality of the herbivory; set at $\alpha = 1$, $g_{i,t}$ will equal the plant's root mass ($r_{i,t}$) in proportion to the total root mass ($R_t$) at time $t$. Above 1, $\alpha$ will increase the preference of the herbivores to disproportionally prefer large root systems (Sonnemann et al. 2013). This parameter is varied from 1 (generalist) to 2 (extremely preferential). If the biomass to be removed from a plant is larger than its total root mass (which may occur, based on the distribution of plant biomasses and $\alpha$), the plant is killed and the overshoot biomass remains in the feeding need ($n_t$), to be removed from other plants.

#### 7.5 Eutrophication

Eutrophication was simulated as an increase in belowground resources (BRes) from the baseline resource rate. Therefore, in IBC-grass a eutrophication intensity of 10 would translate to an increase in belowground resources of 10 BRes for the duration of the experiment. Immediately after the experimental period, the belowground resources return to their pre-eutrophication level for 100 simulation years, for the analysis of the plots' recovery. For this analysis we increased the amount of belowground resources by 50% over their baseline levels (Weiss et al. 2014), increasing from 60 to 90 BRes during the experimental period.

#### 7.6 Seed addition

External seed input takes place in week 20, when all plants disperse their seeds. All PFTs of the regional PFT pool (containing 86 PFTs) are thereby considered as seed source. Seeds are set to the grid by randomly assigning coordinates. The number of seeds added per year per PFT is constant, i.e. if the number of seeds is 1, one seed per PFT per year will be added to the grid.

Seeds added through this pathway must compete to germinate with the seeds that are already present in that grid cell's seed bank. Further, as with all seeds, if their grid cell is overshadowed no seeds will germinate.

Seed addition was used into this version of the simulation to re-incorporate lost species after eutrophication events. Because IBC-grass operates on a "closed systems," any species lost from the community will remain absent unless they are added externally. To mimic long distance seed dispersal from other communities, and thus facilitate the conditions that enable recovery to occur, we use a modicum of seed dispersal per year in the present analysis (10 seeds/PFT/year).

#### 7.7 Trade-offs

Within IBC-grass, submodels (Section 8) are built upon the plants' state variables (e.g. shoot biomass, root biomass) as well as their trait values. These trait values, (e.g. LMR, $m_{max}$, $g_{max}$, and SLA) are critical to understanding the trade-offs within the model, defined by the submodels' equations. Below we further explore these trades-offs by specifically addressing the impacts of each of their component traits on each fundamental equation.

#### 7.7.1 Growth form

##### 7.7.1.1 LMR

LMR defines the growth form of an individual. A high value represents a rosette growth form—low to the ground but almost entirely composed of photosynthetic biomass. Its importance emerges in the equations defining the amount of photosynthetic area available to the individual (A1), asymmetric competition (A3b), conversion of resources to biomass (A4), and herbivory (A7).

In defining photosynthetic biomass (A1), LMR is multiplicative; a low leaf-mass ratio reduces the amount of light gathering biomass. A value of 1 means the entire shoot biomass can gather resources.

In defining asymmetric competition (A3b), the inverse of LMR is used because plants with a lower fraction of leaf tissue (lower LMR) are considered to be higher—more erect—than those with high LMR values—rosettes.

LMR is incorporated into the plant growth submodel (A4) through the utilization of the allometric relation between mass, and growth form (LMR), and SLA. It is incorporated into the loss term, helping to define the maintenance costs of the plant. Within the equation, LMR helps to define the ZOI of the plant. Plants with larger ZOIs (of which LMR is a component) suffer a higher maintenance cost than those with small ZOIs.

LMR factors into grazing in a similar matter as in competition; higher plants are more susceptible to grazing than lower plants. The equation (A7) reflects this by using the inverse of the LMR.

#### 7.7.2 ZOI size — Aboveground herbivory susceptibility

##### 7.7.2.1 SLA

SLA defines the ratio between leaf mass and ZOI area. In other words, a plant that has a high SLA can gather more aboveground resources per mg shoot mass. This relation is incorporated into the competition (A1) and plant growth submodels (A4).

In the competition submodel (A1), SLA is multiplied by the total photosynthetic biomass to produce the ZOI area ($A_{shoot}$). This ZOI area then determines how many grid cells the plant overlaps and can therefore gather resources from.

In the plant growth submodel (A4), SLA helps to define the ZOI of the plant. The maintenance cost (loss term) of the relation includes this ZOI, meaning that plants with larger ZOIs must spend more resources maintaining this photosynthetic biomass than their smaller ZOI competitors.

##### 7.7.2.2 Palat

Palatability is positively correlated to SLA and is integral to calculating the susceptibility of a plant towards grazing. Plants with larger body mass, more erect growth form, and more palatable leaves are more susceptible to grazing.

#### 7.7.3 Growth rate — stress tolerance

##### 7.7.3.1 $g_{max}$

$g_{max}$ reflects maximum number of resources a plant can uptake per week per grid cell. Therefore, plants with a high $g_{max}$ can take up a larger amount of resources per time step than their competitors. It is incorporated into the two versions of competition (A3a, A3b) as well as the intraspecific competition (A3c) and plant growth (A4).

In size-symmetric competition (A3a), the competitive ability of a plant is directly related to its $g_{max}$. In size-asymmetric competition, a plants competitive ability is related to its $g_{max}$, its size, and its growth form (LMR). Larger plants are considered dominant to small plants, and high plants (low LMR) are considered to overshadow low plants (high LMR).

In intraspecific competition (A3c), the number of intraspecific competitors overlapping that same grid cell reduces the $g_{max}$ of a plant on that grid cell.

In plant growth (A4), the $g_{max}$ of a plant helps to define its maintenance cost (loss term). This behaviour operates under the assumption that higher growth rates are generally correlated to higher maintenance costs.

##### 7.7.3.2 $surv_{max}$

$surv_{max}$ is negatively correlated to $g_{max}$. the defines the stress tolerance of a plant. An integer, it represents the maximum number of weeks a plant can survive without meeting its maintenance costs. The probability of mortality linearly increases with each of these weeks.

#### 7.7.4 Maximum plant and seed size — dispersal ability

##### 7.7.4.1 $m_{max}$

The maximum size of the plant is positively correlated to its seed size and incorporated into the plant growth submodel (A4). It is a component of the loss term which defines the maintenance cost of the plant. It is allometrically related to the current size of the plant, in such a way that plants near their $m_{max}$ spend more resources on maintenance costs than do plants far from their $m_{max.}$

##### 7.7.4.2 $m_{seed}$

The mass of a seed is directly correlated to the plant's maximum size ($m_{max}$), and negatively correlated to the mean dispersal distance of the seed ($mean_{disp}$). The value defines the relative probability that the seed will germinate, versus the other seeds on that grid cell, as well as the initial biomass of the newly established plant.

##### 7.7.4.3 $mean_{disp}$

Mean dispersal distance is negatively correlated to maximum plant size and seed size and is integral in defining the dispersal kernel of the plant. Plants with high $mean_{disp}$ can disperse farther than those with lower $mean_{disp}$. Seed dispersal is based on a log-normal distribution, where both the mean and variance parameters are equivalent—this ensures that large seeds have short dispersal distances with little variance. Small seeds, vice versa, disperse far with a longer tail.

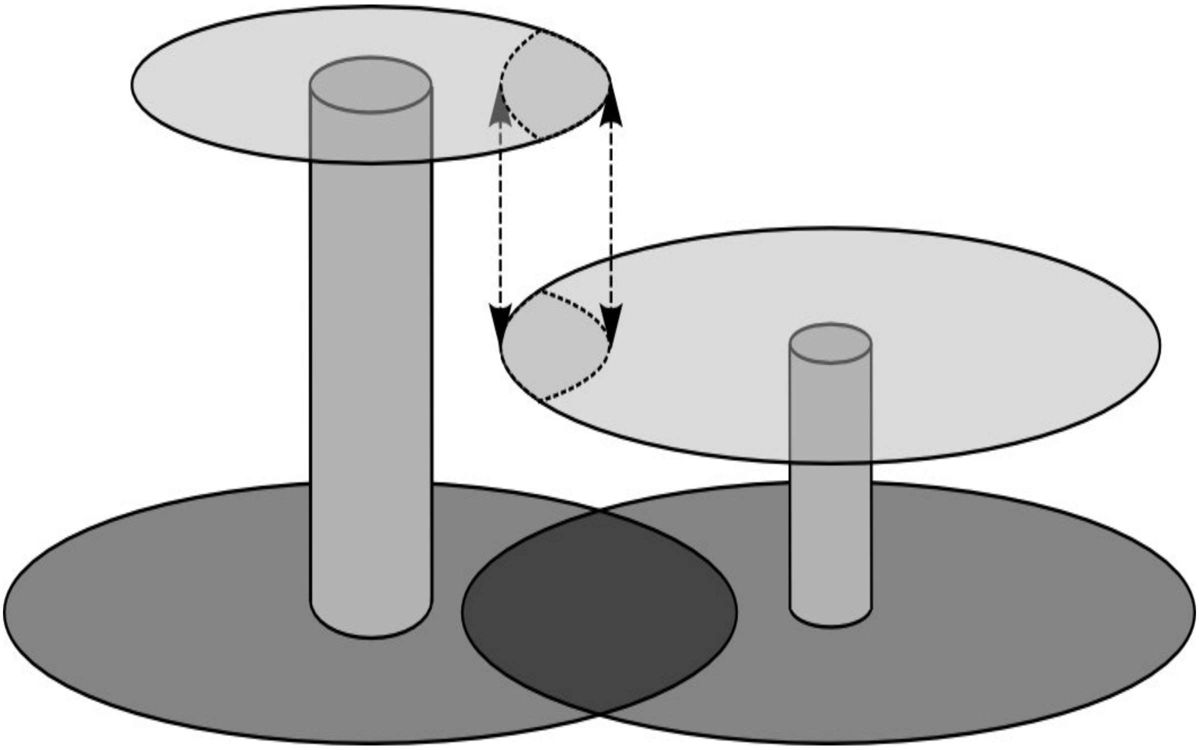

Figure A1. Illustration of the “zone-of-influence” (ZOI) approach including above- and below-ground competition and different shoot geometries. Above- and below-ground “zones-of-influence” are shown as light and dark grey circles, respectively. Stems and support tissue are represented as grey cylinders. Plant individuals compete for resources in the areas of overlap only (arrows indicate the area of above-ground competition). The plant to the left has a lower ratio of leaf mass to shoot mass (LMR) and thus a smaller above-ground ZOI. In return its competitive ability for above-ground resources (light) is higher as it will shade the plant to the right.

Table A1. Model state variables

| State variable | Unit | Description |
|---|---|---|
| **Plants** | | |
| $m_{shoot}$ | mg | vegetative shoot mass (leaves + stems) |
| $m_{root}$ | mg | root mass |
| $m_{repro}$ | mg | reproductive mass (seeds) |
| $w_{stress}$ | weeks | duration of resource stress exposure |
| PFT ID | - | identification number for plant functional type |
| Plant ID | - | Identification number for the individual plant |
| **Seeds** | | |
| $m_{seed}$ | mg | seed mass |
| age | years | time since release from mother plant |
| **Cells** | | |
| $Res_A$ | units/cm² | above-ground resource availability |
| $Res_B$ | units/cm² | below-ground resource availability |

Table A2. PFT parameter values

| Symbol | Description | Unit | Value |
|---|---|---|---|
| **Vegetative traits** | | | |
| $LMR$ | ratio of leaf mass to total shoot mass | mg/mg | ** |
| $SLA$ | above-ground ZOI area per leaf mass | cm²/mg | ** |
| $c_{root}$ | below-ground ZOI area per root mass | cm²/mg | 1.0 |
| g | conversion rate resource units to biomass | mg/resource unit | 0.25 |
| $g_{max}$ | maximal resource utilization per time step and ZOI area (equal for shoot and root*) | resource units/cm²/ week | ** |
| $thr_{res}$ | Threshold fraction of $g_{max}$ considered as resource stress | - | 0.2 |
| $surv_{max}$ | maximal survival time under resource stress exposure | weeks | ** |
| $m_{max}$ | maximum plant mass (equal for shoot and root*) | mg | ** |
| palat | palatability – susceptibility towards grazing | - | ** |
| **Generative traits** | | | |
| $m_{seed}$ | mass of a single seed | mg | ** |
| $mean_{disp}$ | mean of dispersal distance | m | ** |
| $std_{disp}$ | standard deviation of dispersal distance | m | ** |
| $p_{germ}$ | germination probability | - | 0.5 |
| $t_{disp}$ | time of seed dispersal | week of the year | 21 |
| $t_{germ}$ | time of seed germination | week of the year | 1 – 4<br>21 – 25 |

**PFT specific values, see May et al. 2009

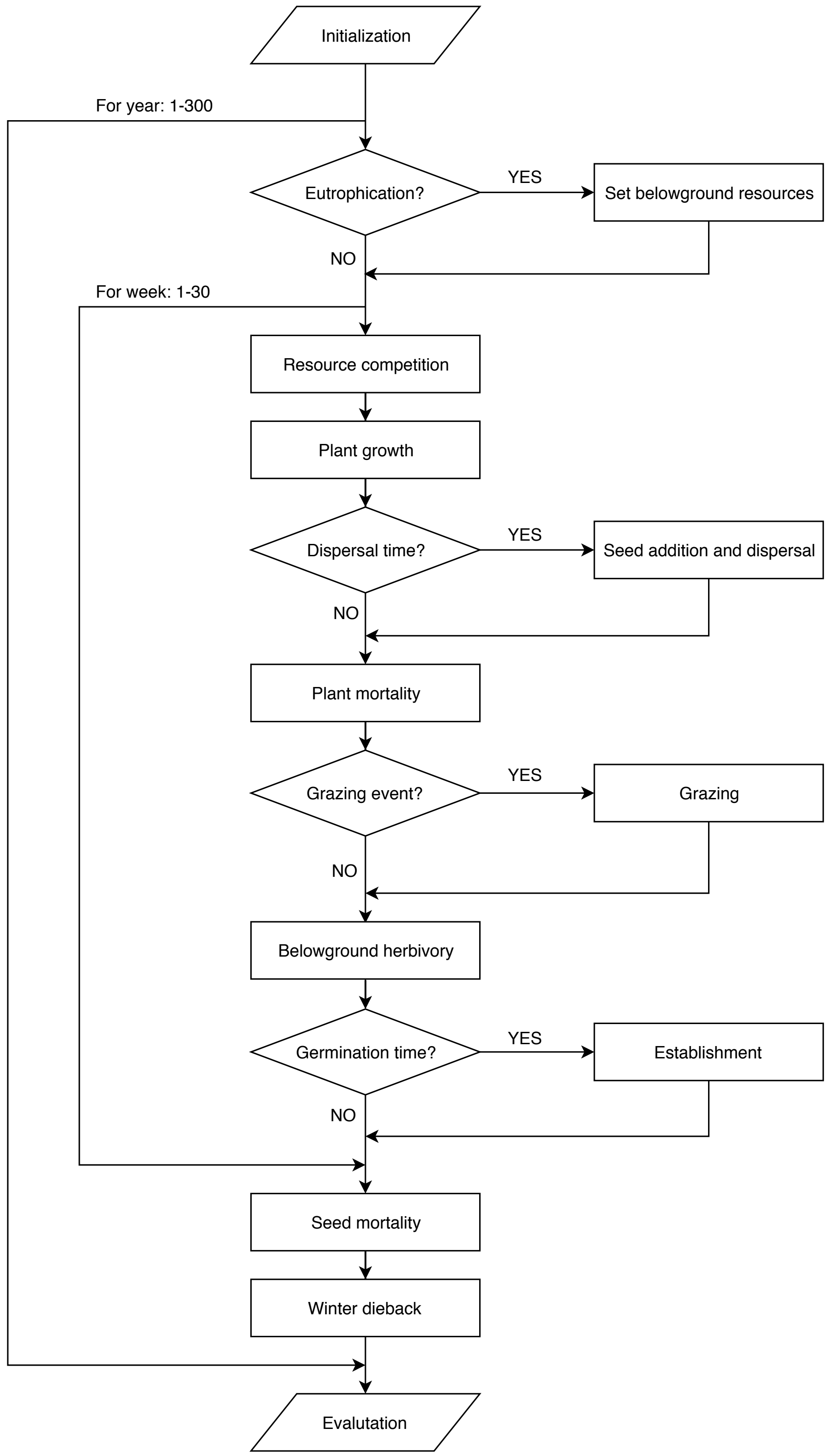


Figure A2. Flow-chart representing process scheduling in the grassland model. Resource competition, plant growth and mortality are executed each week, while seed dispersal and seedling establishment are restricted to certain weeks of the year (see Table A2). Grazing events occur stochastically with a fixed probability per week. Seed mortality and winter dieback are only considered once at the end of each year. State variables of all plants are updated synchronously after each process. Simulations were run for 100 years with 30 weeks vegetation period per year.

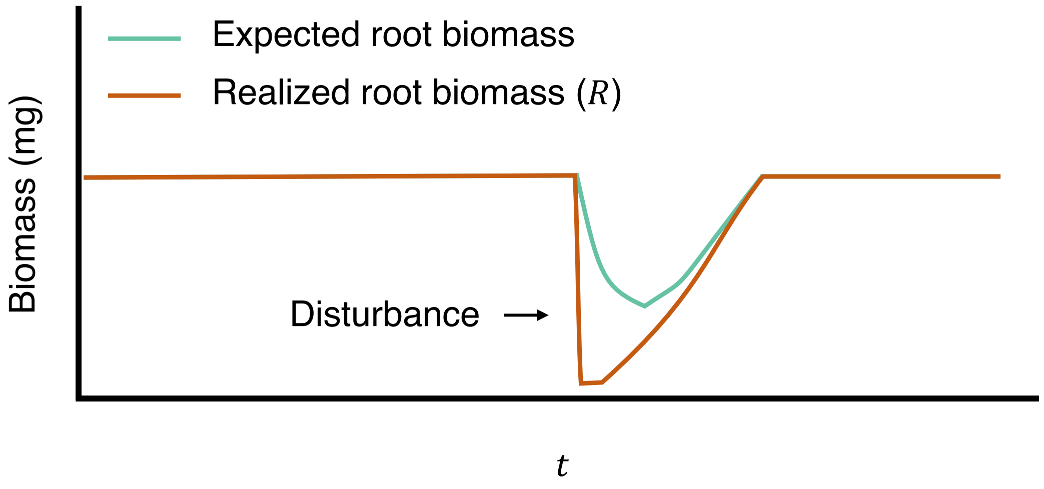


**Figure A3:** A conceptual figure of equation A8. If there is less root mass available than the root herbivores need, the feeding response is triggered in which the herbivores will leave some percentage uneaten. As the realized root biomass gradually increases, the expected root biomass and realized root biomass will begin to equalize.
